# Individual-tree phenology reveals climate-dependent responses to urban thermal heterogeneity across U.S. cities

**DOI:** 10.64898/2026.08.05.740604

**Authors:** Jiali Zhu, Yiluan Song, Juwon Kong, Lin Meng, Michele Peruzzi, Yiwen Zhang, Lei Zhao, Kai Zhu

## Abstract

Rapid urbanization is altering the seasonal life cycles, or phenology, of urban trees. Diverse species composition and heterogeneous urban thermal environments can generate pronounced phenological variation within cities, yet such fine-scale seasonality remains poorly understood. This limits the ability to anticipate localized ecosystem services and disservices, including canopy cooling and pollen exposure. Here, we analyzed individual-tree phenology and their response to temperature across 76 cities in the contiguous United States by integrating municipal street-tree inventories, PlanetScope satellite time series, and 1-km near-surface urban air-temperature data. We quantified intra-city and cross-city phenological variation, estimated species-level associations between phenology and fine-scale urban temperature, and tested whether these associations vary with regional climatic context. Intra-city phenological variation rivaled cross-city variation for 33.3% of species in spring and 53.7% in fall. Temperature-phenology associations were widespread: warmer local urban environments were associated with earlier spring onset in 51.5% of cases and delayed fall senescence in 66.5%, with approximately two-thirds of these responses cascading to longer growing seasons. These associations varied with regional climate, generally showing stronger spring advancement and fall delay in cooler and wetter cities, but weaker or reversed responses in hotter or drier cities. Together, these results suggest that urban tree phenology is shaped by interactions among fine-scale urban temperature, species identity, and regional climate. Accounting for this heterogeneity can improve predictions of urban forest function and inform climate-resilient species selection and management.

**Plain Language Summary:** Street trees within the same city do not all leaf out in spring or lose their leaves in fall at the same time. This seasonal timing, called phenology, affects services such as shade and cooling, as well as disservices such as pollen exposure. We studied individual tree phenology and their temperature response in 76 U.S. cities using street-tree records, satellite observations, and local air-temperature data. We asked how much phenology varies within cities, how it responds to neighborhood-scale temperature differences, and whether these responses depend on regional climate. Variation within a city was often as large as variation among cities, especially in fall. Trees in warmer parts of cities generally leafed out earlier and lost their leaves later, often extending the growing season. However, these responses were stronger in cooler and wetter cities and weaker or sometimes reversed in hotter or drier cities. These findings show that urban tree seasonality depends on local temperature, species identity, and regional climate, which should be considered when selecting and managing trees for future cities.

**Key Points:**

- Substantial within-city phenological variation is not captured by city-level averages.
- Fine-scale temperature shaped phenology and growing-season length, but responses varied across species, cities, and phases.
- Temperature sensitivity depended on regional climate, weakening or reversing in hotter or drier cities.

## 1 Introduction

Rapid urbanization is altering the timing of plant life-cycle events—phenology—in cities, with important implications for urban forestry and localized ecosystem services, including canopy cooling, pollen exposure, carbon uptake, and wildlife interactions (Yang et al., 2023; Zhou, 2022). Evidence for urban phenological shifts includes earlier spring leaf-out and later fall senescence, patterns often attributed to elevated urban temperatures associated with the urban heat island effect (Kabano et al., 2021; Zhang et al., 2006). Most existing evidence, however, stems from urban-rural comparisons or citywide averages, which implicitly treat each city as phenologically uniform (Kabano et al., 2021; Li et al., 2017; Meng et al., 2020; Wang et al., 2019; Zhang et al., 2006). This assumption is unlikely to hold in practice. Cities are spatially heterogeneous mosaics of built surfaces, vegetation, management regimes, and planted species. As a result, individual trees within the same city may experience different local thermal environments and exhibit different phenological timing (Wu et al., 2025). Ignoring this intra-city heterogeneity can obscure the temperature-phenology relationships that operate within cities and limit the use of cities as natural laboratories for understanding ecological responses to climate change (Zhou, 2022). It can also hinder urban-forest management by masking neighborhood-scale variation in ecosystem services and disservices that affect public health and well-being (Crawford et al., 2024; Katz et al., 2019; Song et al., 2025).

Increasing evidence shows that phenological timing can vary markedly among individual trees as a function of the local environments they experience (Christmann et al., 2023; Wohlfahrt et al., 2019; Wu et al., 2024; Zellweger et al., 2020). In cities, two sources of variation are likely to be especially important: fine-scale climate variation and species diversity. First, individual trees experience substantial intra-city variation in air temperature, a major environmental cue for plant phenology (Peñuelas et al., 2009; Piao et al., 2019; Wohlfahrt et al., 2019). This thermal heterogeneity arises from spatial variation in urban form, surface materials, vegetation cover, and anthropogenic heat emissions (Stewart and Oke, 2012). Reported intra-city seasonal temperature differences can reach up to 5–6 ^◦^C, comparable to or greater than many urban-rural contrasts (Cao et al., 2021; Hart and Sailor, 2009; Johnson et al., 2020; Kong et al., 2025; Kousis et al., 2021; Naserikia et al., 2023; Venter et al., 2020). Second, urban trees are composed of diverse assemblages of intentionally planted species with distinct phenological schedules and environmental sensitivities (Gaš parović et al., 2023; Parece and Campbell, 2018). Because multiple species are often planted across the same neighborhoods and streets, species composition can amplify observed phenological heterogeneity and motivate analyses that explicitly link individual-level phenology to taxonomic identity.

Although single-city studies have documented intra-urban variation in tree phenology, whether these patterns generalize across broader climatic contexts remains unclear (Crawford et al., 2024; Katz et al., 2019; Parece and Campbell, 2018; Zipper et al., 2016; Wu et al., 2025). Regional climate may modify temperature-phenology relationships through chilling limitation, heat stress, moisture constraints, and differences in species composition or provenance (Chen et al., 2020; Chuine et al., 2010; Jochner and Menzel, 2015; Fu et al., 2015; Li et al., 2019; Zhou, 2022). However, it remains poorly understood how these broader climatic constraints interact with fine-scale intra-urban thermal heterogeneity. The contiguous United States provides a useful setting for addressing this issue because it spans large gradients in climate, urban form, and tree species composition. At the same time, the availability of municipal tree inventories, high-resolution satellite imagery, and gridded urban air-temperature products enables analysis of phenology at the level of individual trees across many cities.

Here, we examine individual-tree phenology across 76 cities in the contiguous United States by integrating municipal street-tree inventories, PlanetScope satellite time series, and 1-km near-surface urban air-temperature data. We address three questions. First, how large is intra-city variation in tree phenology relative to variation among cities? We retrieve start-of-season (SOS) and end-of-season (EOS) metrics for more than 1.6 million inventoried deciduous street trees from 345 species and partition phenological variation into intra-city, cross-city, and interannual components. We expect substantial intra-city variation because individual trees experience different local environments and urban tree communities have diverse species. Second, how is individual-tree phenology associated with fine-scale intra-city temperature variation? For each species within each city, we estimate the association between phenological timing and local pre-season air temperature while accounting for precipitation, radiation, spatial autocorrelation, and interannual variability. We expect warmer local environments to be associated with earlier SOS, later EOS, and consequently longer growing-season-length (GSL) for many species, while also expecting substantial variation in magnitude and direction among species–city combinations. Third, do these temperature-phenology associations vary with regional climatic context? We test whether estimated temperature sensitivities vary with city-level mean annual temperature and precipitation. We expect apparent temperature sensitivities to weaken or reverse in hotter or drier cities, where chilling limitation, heat stress, or moisture constraints may limit phenological advancement or growing-season extension.

## 2 Materials and methods

### 2.1 Data sources

#### 2.1.1 Urban tree inventories

To obtain the exact locations and species identity of individual urban trees, we compiled municipal street-tree inventories across cities in the contiguous United States (Table S2). We first standardized two published inventory datasets: an assembly by McCoy et al. (2022) and a database on OpenTrees.org. The dataset compiled by McCoy et al., was assembled through Google searches and connections with relevant state officials, limited to the higher-populated U.S. cities. This dataset provides detailed information on tree location, species, and health status. We retained only trees located within urban boundaries defined by TIGER/Line Files from the U.S. Census Bureau, excluding individuals with “dying” or “dead” conditions as well as duplicate records. This yielded inventories for 40 cities. OpenTrees.org is a public platform that aggregates tree inventories from municipalities worldwide. We selected 10 cities from this database with clearly documented government data sources. For these cities, we standardized tree coordinates, removed duplicate records, and filtered trees by urban boundaries. To improve the representation of urban diversity, we further added 26 additional cities considering their climatic, demographic, and built-environment characteristics (SI Text S1). The selected cities were broadly representative of urban conditions across the contiguous United States (Fig. S1).

To harmonize taxonomy, we matched species names to the GBIF backbone taxonomy using the R package rgbif (Chamberlain et al., 2024). For inventories reporting only common names, we converted them to scientific names with the R package taxize (Chamberlain et al., 2025). We restricted our analysis to deciduous trees because their clear seasonal leaf phenology is readily detectable using remote sensing. Species were assigned to a categorical leaf phenology type (TraitID = 37) based on the most frequently observed classification in the TRY database (Kattge et al., 2011). To ensure adequate sample sizes, we excluded species with fewer than 30 individuals per city. For species with more than 2,000 individuals in a city, we randomly downsampled to 2,000 trees (Psutka and Psutka, 2019). After processing, the final dataset included 1,663,951 street trees from 345 species across 76 cities in the contiguous United States (Fig. S1, Fig. S13).

#### 2.1.2 Individual tree phenology

We extracted SOS, EOS, and GSL phenological metrics for individual trees from PlanetScope-derived enhanced vegetation index (EVI) time series, following the general workflow of Song et al. (2025). The daily-temporal and 3-m spatial resolution satellite imagery from PlanetScope is particularly well positioned for high-resolution vegetation monitoring across broad spatial extents. Previous studies have shown the reliability of using PlanetScope imagery to detect phenological events of individual trees (Moon et al., 2021; Song et al., 2025; Zhao et al., 2022), given that many canopy trees have crown diameters exceeding PlanetScope’s spatial resolution (Dixon et al., 2025; Liu et al., 2023; Zhang et al., 2025). This advantage is even more pronounced in urban settings, where the sparse distribution of urban trees against contrasting backgrounds enhances the detectability of their temporal phenological signals (Liu et al., 2023; Neyns et al., 2024).

We retrieved PlanetScope imagery (ortho analytic 4b sr) from 2017 to 2023 for all trees via the Planet API using the R package BatchPlanet (Song et al., 2026). Data were harmonized to Sentinel-2 for consistency (Kington and Collison, 2022), and only daytime images (sun elevation > 0) were used. Reflectances in the red, green, blue, and near-infrared bands were extracted at tree coordinates for EVI calculation, with UDM2 masks applied to retain only clear pixels (no snow, ice, shadow, haze, or cloud, and classification confidence ≥ 80%). For days with multiple acquisitions, mean reflectances were computed.

To derive tree-level phenological metrics from PlanetScope imagery, we (i) computed the enhanced vegetation index (EVI) and removed implausible values outside the range of 0–1 (Huete et al., 2002), (ii) extended and smoothed annual time series with weighted Whittaker smoothing to capture full green-up and green-down cycles (Kong et al., 2019), (iii) identified and selected time series of EVI with significant seasonality using piecewise regression models, (iv) extracted the timing of SOS and EOS as the day of year when their growing season EVI curves first crossed the 50% green-updown threshold, with 40% and 60% thresholds evaluated in sensitivity analyses (SI Text S6); and (v) calculated GSL as the interval between SOS and EOS.

We validated PlanetScope-derived SOS and EOS against ground observations from the USA National Phenology Network (NPN, Rosemartin et al. (2018)) and National Ecological Observatory Network (NEON, Elmendorf et al. (2016)) (SI Text S3). To align field-based observations with PlanetScope-derived phenology metrics, ground-based SOS and EOS were calculated from individual-level phenometrics (Xin et al., 2020; Zhao et al., 2022). The validation dataset included 734 individual deciduous trees representing 123 species across 92 USA-NPN locations in 19 cities from 2016 to 2024, and 987 individual trees representing 37 species across 19 NEON sites from 2018 to 2022. PlanetScope-derived phenology metrics were generally consistent with the corresponding ground-based observations, with stronger agreement for SOS than for EOS, as indicated by higher correlation coefficients and lower RMSEs for SOS (Fig. S2). These results suggest that PlanetScope time series capture meaningful phenological signals and support their application in urban areas.

#### 2.1.3 Climate datasets

To characterize intra-city climate conditions, we used the Urban High-resolution Air Temperature dataset, hereafter U-HAT (Zhang et al., 2026). U-HAT provides daily near-surface, 2-m air temperature for 384 cities in the contiguous United States from 2013 to 2023 at 1-km spatial resolution. This novel dataset uses a transfer learning framework that integrates satellite land surface temperature, urban surface properties, atmospheric forcings, and urban station observations to estimate near-surface air temperature in cities. Previous evaluations showed that U-HAT better captures urban heat island effects and fine-scale thermal variation than alternative temperature products (Zhang et al., 2026). Specifically, gridded weather datasets without urban correction (e.g., Daymet) systematically underestimated urban temperatures, while observations from urban personal weather stations (e.g., Weather Underground) were prone to outliers and data quality issues (SI Text S3, Fig. S4).

For each tree and year, we extracted preseason air temperature from U-HAT. For SOS analyses, preseason temperature was defined as the mean winter temperature from December 1 to February 28 or 29. For EOS analyses, preseason temperature was defined as the mean summer temperature from June 1 to August 31. For GSL analyses, both winter and summer preseason temperature windows were considered separately. We used “fine-scale urban temperature” to refer to intra-city variation in 1-km near-surface air temperature, while recognizing that tree-scale thermal environments can vary at finer spatial scales. To account for other climatic influences on phenology, we also extracted preseason precipitation and preseason shortwave radiation from Daymet version 4 at 1-km spatial resolution (Thornton et al., 2021). These variables were calculated over the same preseason windows used for temperature.

We used mean annual temperature and mean annual precipitation to characterize the city-level baseline climate. These two bioclimatic variables were obtained from CHELSA v.1.2 climatologies at 30 arcsec spatial resolution (Karger et al., 2017). For each city, we extracted values at sampled tree locations and averaged them across trees to represent the baseline climatic conditions experienced by the inventoried street-tree population in that city.

### 2.2 Statistical analyses

#### 2.2.1 Comparing intra-city and cross-city variation in phenology

To compare the relative contributions of different sources of phenological variation, we partitioned variance in SOS and EOS into cross-city, intra-city, and cross-year components. This analysis was conducted separately for each species that occurred in at least two cities. We used a mixed-effects model analogous to a variance-component ANOVA, while allowing nested observations across cities, individuals, and years.

For each species *k*, we modeled phenological timing as shown in Eq. 1. Because the model was fitted separately for each species, the species index *k* was omitted from all terms for notational simplicity:

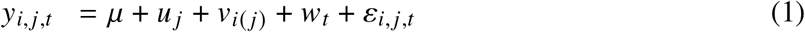

where *y_i_*, *_j_*_,*t*_ is the SOS or EOS of tree *i* in city *j* and year *t*. The parameter *μ* represents the species-specific intercept, or the overall mean phenological timing for species, averaged across cities, individual trees, and years. *u _j_* is the cross-city random effect for species. *v_i_*_(_ *_j_*_)_ is the intra-city (cross-individual) random effect, with individuals nested within cities. *w_t_* is the interannual random effect, and *ε_i_*, *_j_*_,*t*_ is the residual error (Eq. 1).

The random effects and residual error were specified as:

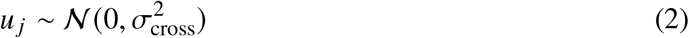

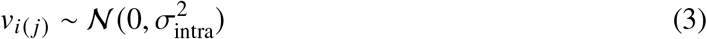

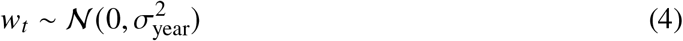

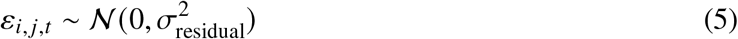

After fitting the model, we extracted the estimated variance components and calculated total variance (Eq. 6). To compare intra-city and cross-city contributions directly, we calculated the proportional contributions (*C*) of cross-city and intra-city variance components relative to their combined variance (Eq. 7). We used these proportions to evaluate whether phenological variation within cities was comparable to or greater than variation among cities for each species.

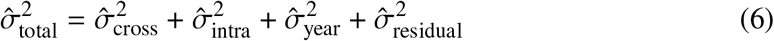

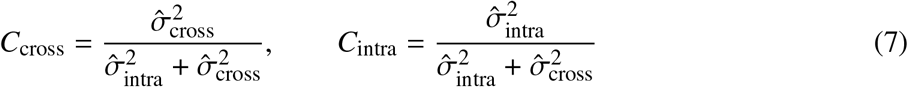

#### 2.2.2 Estimating phenological associations with fine-scale urban temperature

To estimate how individual-tree phenology was associated with fine-scale intra-city temperature variation (Question 2), we fitted spatial statistical models separately for each species–city combination. Spatial models extend standard regression by accounting for spatial autocorrelation among nearby observations, with correlation typically declining as geographic distance increases (Banerjee et al., 2025; Finley et al., 2007). This structure was applied because neighboring trees are more likely to share similar local environments, planting conditions, and management regimes, which can lead to similar phenological patterns. Accounting for this spatial structure helps isolate phenological responses to temperature variation rather than confounding them with unmodeled spatial dependence.

For each species–city combination (*k*, *j*), we modeled individual-tree phenology as a function of local climatic conditions while accounting for spatial and temporal dependence (Eq. 8). Because the model was fitted separately for each combination, the species and city indices *k* and *j* were omitted from all terms for notational simplicity:

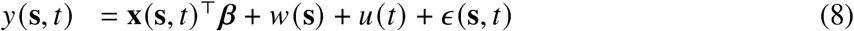

where *y*(**s**, *t*) is the observed annual phenological timing, measured as the day of year (DOY) of SOS, EOS, or GSL for a tree at geographic coordinate **s** in year *t*. **x**(**s**, *t*) is the vector of fine-scale climate predictors. ***β*** is the vector of regression coefficients. *w*(**s**) is the spatial random effect, and *u*(*t*) is the temporal random effect. *ε* (**s**, *t*) is the residual term.

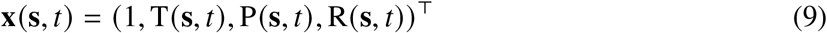

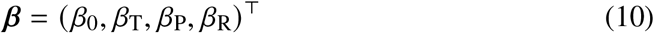

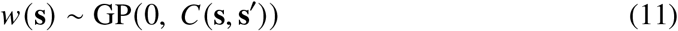

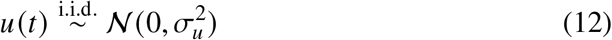

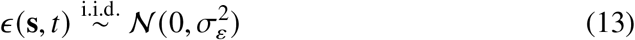

Here, T(**s**, *t*), P(**s**, *t*), R(**s**, *t*) represent preseason temperature, precipitation, and shortwave radiation, respectively. The coefficient for the average temperature, *β*_T_, is the phenological temperature sensitivity, which is the main interest in this study. The spatial random effect, *w*(**s**), follows a Gaussian process with a Matérn covariance structure, *C*(**s**, **s**′). The year random effect *u*(*t*) is modeled as an independent normal deviation with variance σ^2^_*u*_, capturing annual shifts in phenological timing shared across trees. The residual error term *ε* (**s**, *t*) is modeled as independent and normally distributed, with ϵ(**s**, *t*) representing the residual variance.

We used a Bayesian hierarchical modeling framework to jointly estimate fixed climatic effects and latent spatial and temporal random effects, thereby formulating the model as a hierarchical spatial model. This approach also allowed uncertainty in the temperature sensitivity estimates to be propagated into subsequent analyses. Weakly informative priors were assigned to the regression coefficients (Eq. 14).

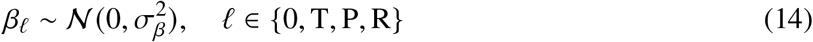

We implemented model fitting using the INLA package in R, which approximates the continuous spatial Gaussian field through the SPDE framework, enabling more efficient Bayesian inference (Rue et al., 2009).

To evaluate the robustness of our results to different model fitting frameworks, we also fitted comparable spatial models using two alternative approaches. First, we used the spBayes package in R, employing a low-rank predictive process approach to reduce computational complexity (Finley et al., 2015, 2007). Second, we fitted frequentist spatial linear models using the spmodel package in R, which estimates spatial covariance parameters using likelihood-based optimization (Dumelle et al., 2023). Results reported in the main text were obtained using the INLA approach, whereas results from spBayes and spmodel were presented in SI Texts S4 and S5.

#### 2.2.3 Testing whether phenological sensitivities varied with regional climate

To evaluate whether phenological temperature sensitivities varied with regional climatic context (Question 3), we used the species–city temperature coefficients estimated from the spatial models described above as response variables. Because these coefficients were estimated with uncertainty, we propagated this uncertainty using a parametric resampling approach based on the approximate posterior distributions returned by INLA. Specifically, for each species–city coefficient, we sampled values from a normal distribution parameterized by its INLA posterior mean and standard deviation. Each draw produced one iteration-specific dataset of temperature sensitivities. For each iteration *r*, we modeled the sampled phenological sensitivity in Eq. 15:

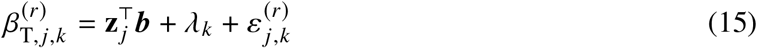

where β^(*r*)^_T,*j,k*_ represents the *r*-th posterior draw of the temperature coefficient estimated in Section 2.2.2 for species *k* in city *j* at iteration *r*. This coefficient represents the temperature sensitivity of phenological timing, SOS or EOS. **z** *_j_* is the vector of regional climatic predictors. ***b*** is the vector of regression coefficients. *λ_k_* represents the species-level random effect, an ε^(*r*)^_*j,k*_ is the residual term.

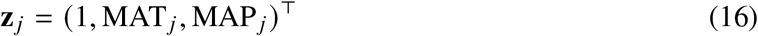

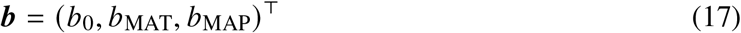

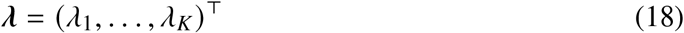

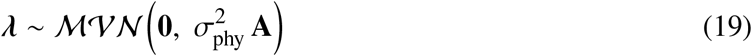

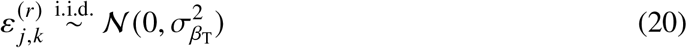

The predictor vector **z** *_j_* includes an intercept, mean annual temperature (MAT*_j_*), and mean annual precipitation (MAP*_j_*) for city *j* . ***b*** = (*b*_0_, *b*_MAT_, *b*_MAP_)^⊤^ are the regression coefficients for the intercept and covariates. The species-level random effect, *λ_k_*, accounts for the possibility that closely related species may show similar phenological sensitivities due to shared evolutionary history (Li et al., 2020; Losos, 1996). Collectively, the species-level random effects are assumed to follow a multivariate normal distribution with covariance proportional to the phylogenetic relatedness matrix **A**. ε^(*r*)^_*j,k*_ is modeled as independent and normally distributed, with σ^2^_β*T*_ representing the residual variance.

We also used a Bayesian approach to jointly estimate climatic effects and phylogenetically structured species-level variation, allowing us to account for species non-independence while quantifying uncertainty in climate effects on phenological temperature sensitivity. Weakly informative priors were assigned to the regression coefficients ***b*** (Eq. 21).

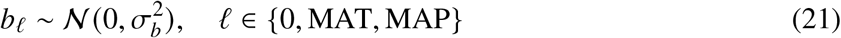

Final inference was obtained by aggregating parameter estimates across 1,000 iterations.

For visualization, we also fitted separate univariate models between temperature sensitivity and each climatic variable for common species. These visualization models were not used as the primary basis for inference but helped illustrate bivariate patterns between temperature sensitivity and regional climate.

## 3 Results

### 3.1 Widespread intra-city variation in individual-tree phenology

Intra-city phenological variation was substantial for many species. For SOS, intra-city variation exceeded cross-city variation for 85 of 254 species. For EOS, this pattern was observed in 137 of 255 species. We illustrated these patterns using *Pistacia chinensis* (Chinese pistache) for SOS and *Quercus alba* (White oak) for EOS, two species for which intra-city differences accounted for a larger share of phenological variation.

For *Pistacia chinensis*, spring phenology varied markedly among individual trees within cities (Fig. 1a). Although the interquartile range of city-level median SOS across cities was relatively narrow, spanning DOY 92–107, individual cities often showed much broader phenological spread. For example, three cities located in different regions all showed broad intra-city variation: Los Angeles, CA, which had the earliest spring onset, showed a 64-day intra-city interquartile range, while Portland, OR and New Orleans, LA had similar city-level median SOS values but still showed intra-city interquartile ranges of about 31–32 days. Variance partitioning showed that intra-city differences accounted for 54.48% of the combined intra-city and cross-city SOS variance in *Pistacia chinensis* (red point in Fig. 1b).

**Figure 1:**
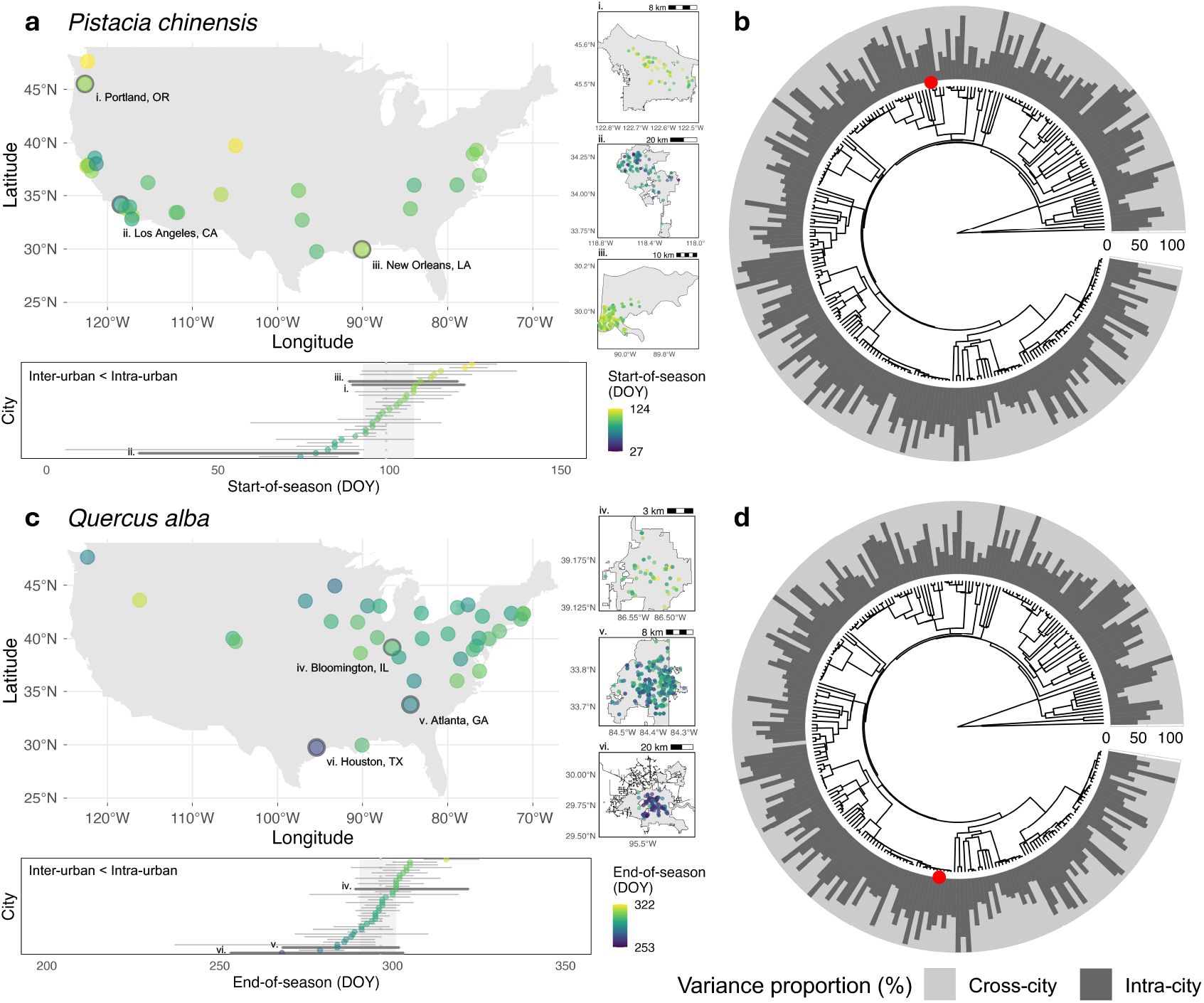
Phenological distributions and variance partitioning of intra-city and cross-city components. (a) Distribution of SOS for *Pistacia chinensis* in 2022. (b) Proportional contributions of intra-city and cross-city components to variation in SOS across species. (c) Distribution of EOS for *Quercus alba* in 2022. (d) Proportional contributions of intra-city and cross-city components to variation in EOS. In (a) and (c), points on the maps indicate city-level mean phenology, and insets show individual-tree variation within three selected cities. Horizontal interval plots below each map show intra-city phenological distributions, with points representing city means and horizontal lines indicating interquartile ranges. Dark lines highlight the distributions of the three example cities shown on the maps and insets, while the gray shaded regions indicate the range of city-level means. In (b) and (d), each bar associated with the phylogenetic tree represents one species, and the red spots highlight the two example species shown in (a) and (c).

Similarly, *Quercus alba* showed broad intra-city variation in fall phenology (Fig. 1c). The interquartile range of city-level median EOS spanned only 10.5 days across cities (DOY 290.5–301), three cities of different sizes—Houston, TX, Atlanta, GA, and Bloomington, IL—showed much broader intra-city variation, with interquartile ranges reaching 33–50 days. Variance partitioning showed that intra-city differences accounted for 65.38% of the combined intra-city and cross-city EOS variance in *Quercus alba* (red point in Fig. 1d).

Across all species, phenological observations spanned a broad seasonal range (Fig. S14), with most SOS observations occurring between March and May and most EOS observations occurring between August and November. Variance partitioning showed large interspecific differences in the relative importance of intra-city variation (Fig. 1b, d). For SOS, the proportion of intra-city variance ranged from 4.21% to 73.73% across species. For EOS, it ranged from 0.08% to 63.73%. In spring, 85 species were dominated by intra-city variance and 169 species were dominated by cross-city variance. In fall, 137 species were dominated by intra-city variance and 118 species were dominated by cross-city variance. Although variance components were visualized along the phylogenetic tree, high and low proportions of intra-city variance were distributed across multiple clades, with no obvious clustering within particular evolutionary lineages (Fig. 1b, d).

### 3.2 Individual-tree phenology associated with fine-scale urban temperature variation

Within cities, phenological timing was associated with fine-scale air temperature variation for many species–city combinations. Across 3,121 species–city combinations for SOS, 51.5% of species– city combinations had negative temperature coefficients (30.7% significant), indicating earlier leaf-out under warmer climate conditions (Fig. 2a). Among the significant negative responses, the interquartile range of temperature sensitivity was -1.54 to -0.53 days ^◦^C^−1^, indicating an approximately 0.53–1.54 day advance in SOS per 1 ^◦^C warming. Moreover, among the 1,607 species–city combinations showing earlier SOS under warmer winters, 61.4% also exhibited longer GSL. Across these combinations, the mean GSL sensitivity was 0.48 days ^◦^C^−1^, corresponding to an average extension of 0.48 days in GSL per 1 ^◦^C increase in winter temperature (Fig. 2c).

**Figure 2:**
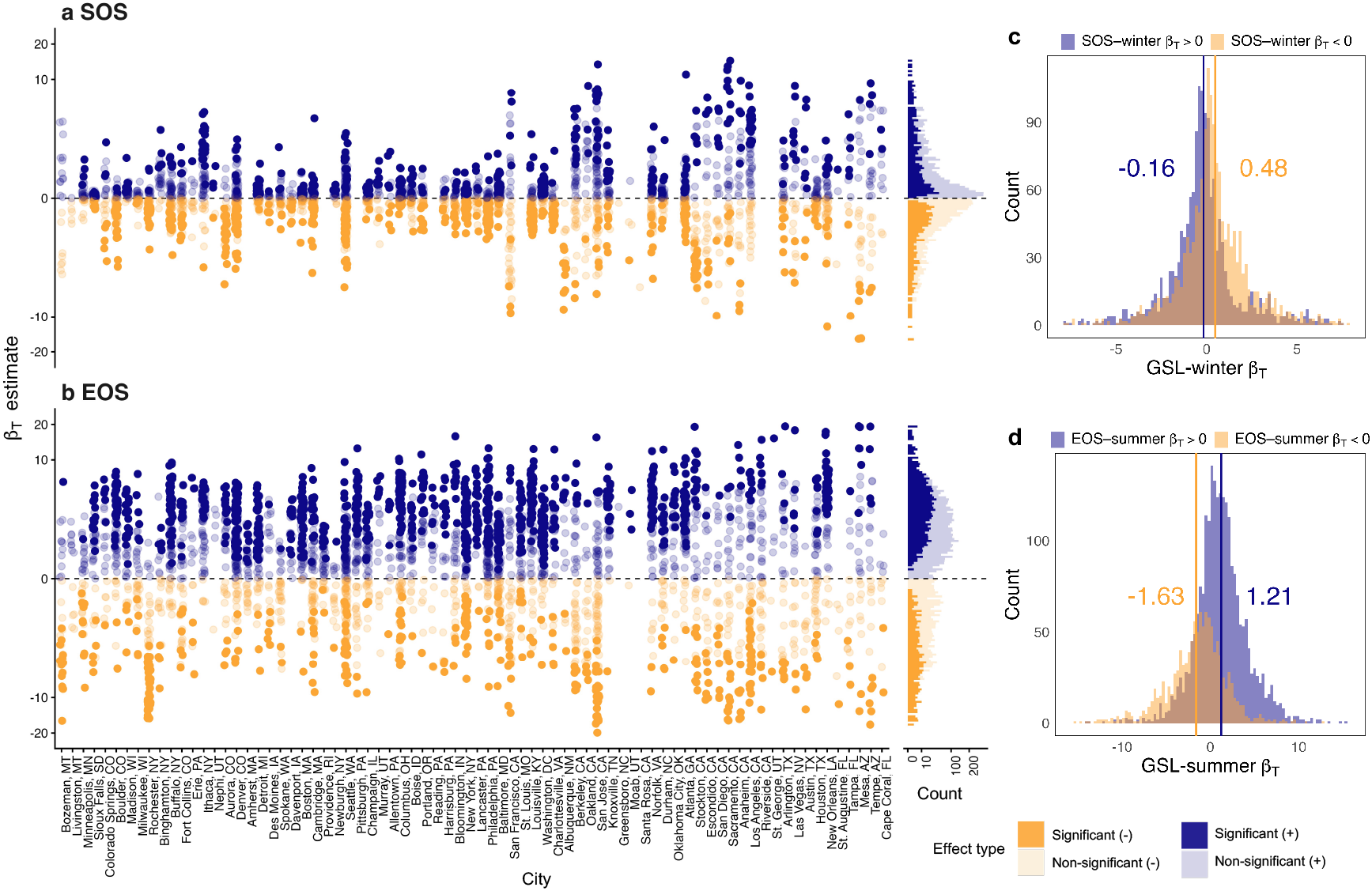
Phenological associations with fine-scale intra-city temperature variation across species and cities. (a, b) Estimated temperature coefficients for SOS (a) and EOS (b) across all species–city combinations. Each point represents one species–city combination. Colors indicate the direction of the estimated temperature–phenology association, and color intensity indicates statistical significance. Cities along the x-axis are ordered by mean annual temperature (bio1). The marginal stacked histograms on the right show the frequency distributions of the estimated temperature coefficients. (c, d) Distributions of GSL coefficients for winter temperature (c) and summer temperature (d), grouped according to the direction of the corresponding SOS or EOS temperature response. Vertical lines indicate the mean GSL coefficient within each response group.

In contrast, for EOS, temperature coefficients were more consistently positive. Across 3,486 species–city combinations, 66.5% had positive temperature coefficients, indicating later fall senescence in warmer local urban environments (Fig. 2b). Among these positive effects, 47.4% were significant, with the interquartile range of 2.31 to 5.33 days ^◦^C^−1^, indicating an approximately 2.3–5.3 day delay in EOS per 1 ^◦^C warming. Similarly, among the 2,318 species–city combinations showing later EOS under warmer summers, 68.4% also exhibited longer GSL, with a mean extension of 1.21 days per 1 ^◦^C increase in summer temperature (Fig. 2d).

Despite these dominant patterns, phenology-temperature associations varied strongly among species and cities in both direction and magnitude. This heterogeneity was especially pronounced for SOS, where responses ranged from strong advances under warmer temperatures to delayed responses in 48.5% of species–city combinations (Fig. 2a). EOS responses were more consistently positive overall, but their magnitude still varied substantially among species–city combinations (Fig. 2b).

Representative species–city examples illustrated this heterogeneity in both spring and fall phenology (Fig. 3). For SOS, a 1 ^◦^C increase in preseason air temperature was associated with a 0.19-day advance for *Acer rubrum* (Red maple) in New York City, NY, and a 4.46-day advance for *Betula pendula* (Silver birch) in Stockton, CA. In contrast, warming delayed spring phenology in other cases, by 2.93 days for *Acer rubrum* in Tampa, FL and 4.63 days for *Betula pendula* in Los Angeles, CA (Fig. 3a–d). For EOS, a 1 ^◦^C increase in preseason air temperature was associated with a 2.43-day delay for *Acer rubrum* in Washington, DC, but a 3.11-day advance in San Francisco, CA. Similarly, *Liquidambar styraciflua* (Sweetgum) showed delayed fall by 2.36 days per ^◦^C in Seattle but advanced fall by 4.36 days per ^◦^C in Sacramento, CA (Fig. 3e–h).

**Figure 3:**
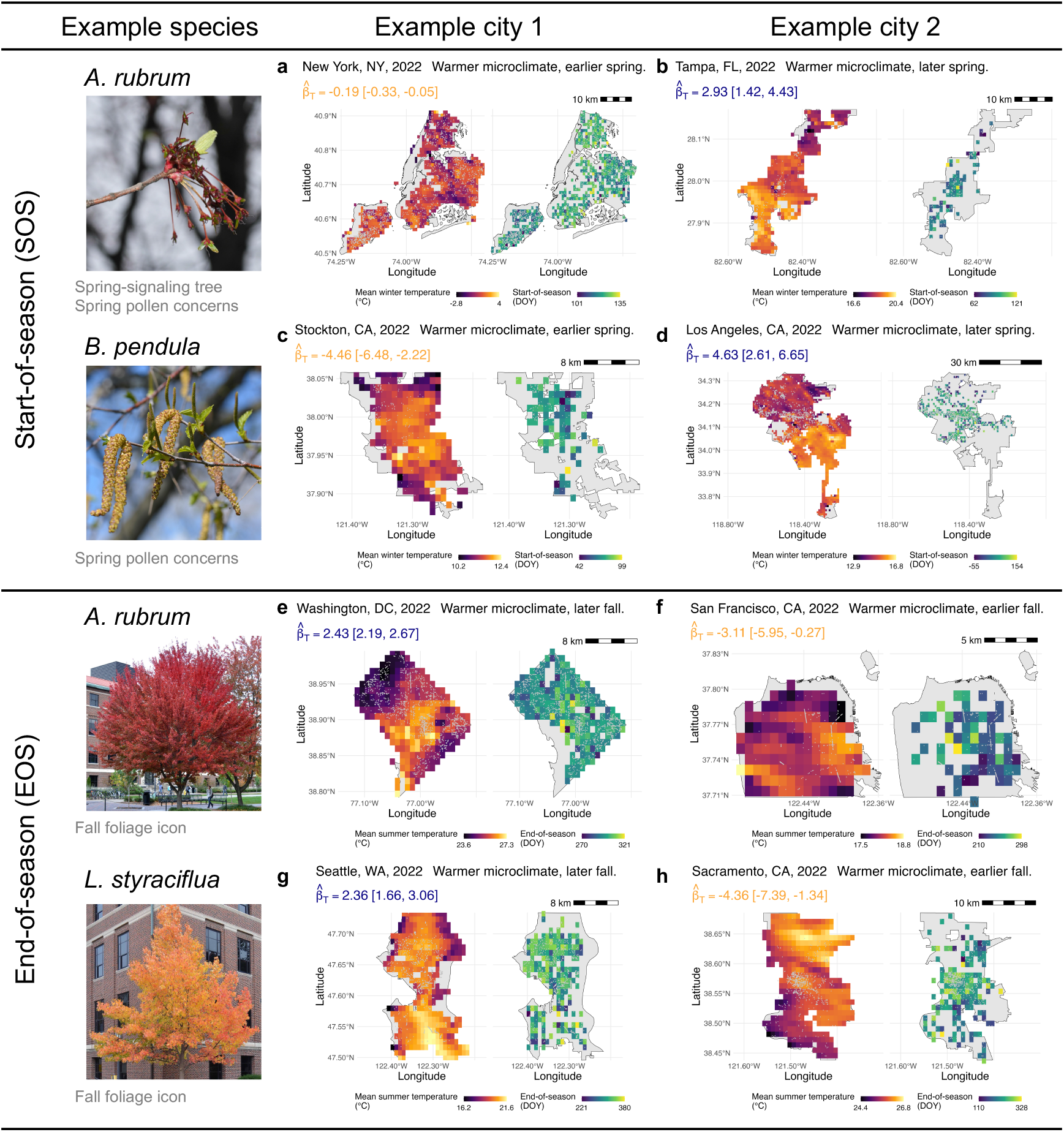
Spatial patterns of mean seasonal temperature and the phenological metric for representative species in eight example cities in 2022. (a) SOS of *Acer rubrum* in New York, NY. (b) SOS of *Acer rubrum* in Tampa, FL. (c) SOS of *Betula pendula* in Stockton, CA. (d) SOS of *Betula pendula* in Los Angeles, CA. (e) EOS of *Acer rubrum* in Washington, DC. (f) EOS of *Acer rubrum* in San Francisco, CA. (g) EOS of *Liquidambar styraciflua* in Seattle, WA. (h) EOS of *Liquidambar styraciflua* in Sacramento, CA. In each panel, the left map shows mean preseason temperature, and the right map shows the aggregated phenological metric at the same pixel resolution as the temperature map. The gray points represent individual tree locations. Photo source: The Purdue Arboretum Explorer (https://www.arboretum.purdue.edu/explorer/)

### 3.3 Temperature–phenology associations varied with regional climatic context

Estimated phenological temperature sensitivities to temperature varied across cities in the contiguous U.S. (Fig. 4a, b). For SOS, 36 of 68 cities showed negative city-level mean temperature sensitivities, with these negative sensitivities concentrated across much of the central and eastern U.S. By contrast, several warmer or drier cities, particularly in California, Texas, and Florida, showed positive sensitivities, indicating delayed spring onset under warmer climatic conditions (Fig. 4a). Specifically, a 1 ^◦^C increase in climatic temperature was associated with delayed SOS by 1.79 days in Los Angeles, CA, 1.66 days in Cape Coral, FL, 1.64 days in Sacramento, CA, and 1.08 days in San Jose, CA. Across cities, the mean intra-city standard deviation was 1.42 days ^◦^C^−1^, with an interquartile range of 0.48–2.30 days ^◦^C^−1^. In other words, within a typical city, species differed in their spring temperature sensitivities by roughly 1–2 days per 1 ^◦^C warming.

**Figure 4:**
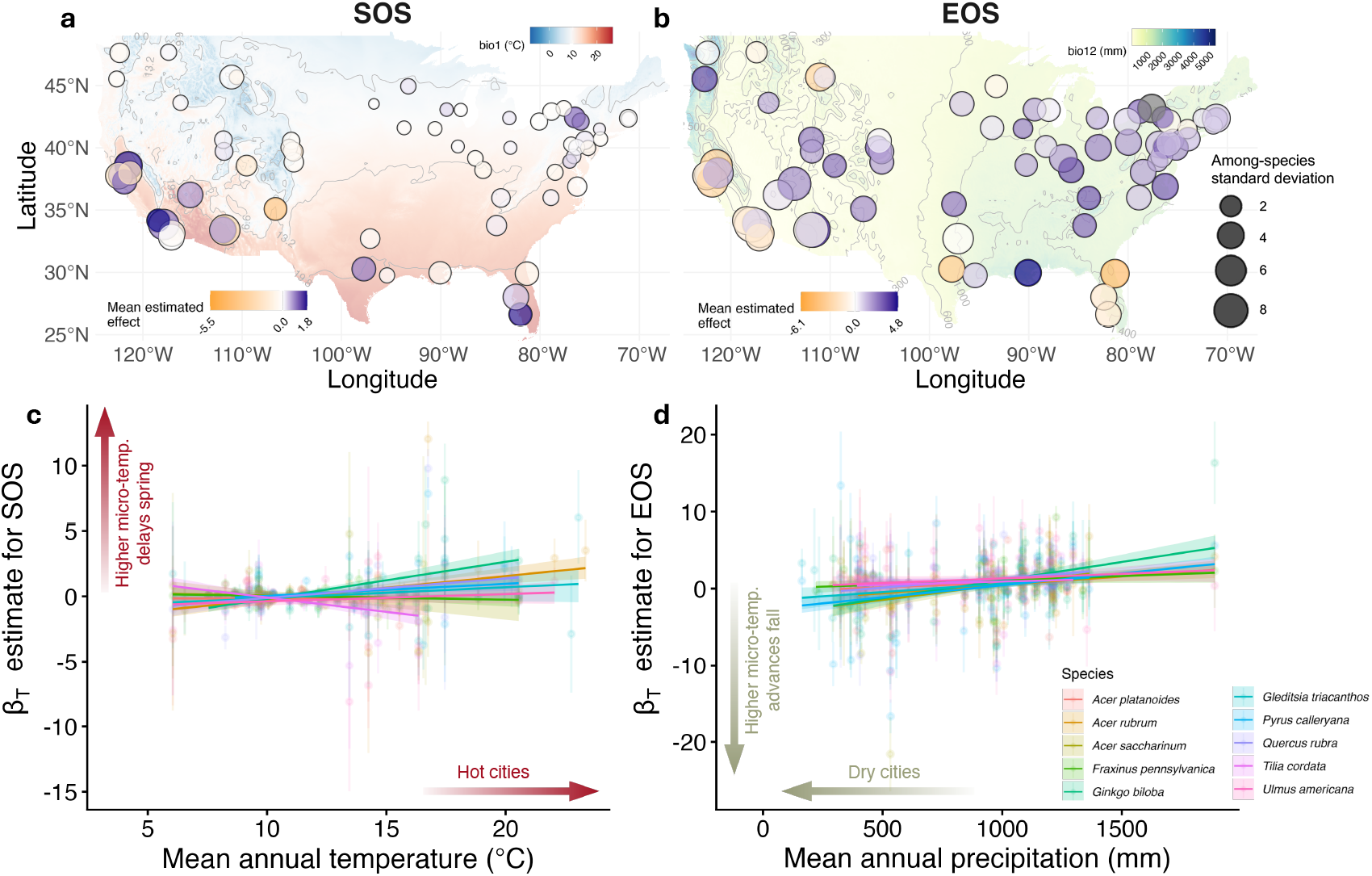
Phenological temperature sensitivities across U.S. cities and their relationships with regional climate. (a, b) Spatial distribution of city-level mean temperature sensitivity for SOS (a) and EOS (b), averaged across species present in each city. Colors represent the magnitude of the average phenological temperature effect, and point sizes indicate the variance of species-specific temperature sensitivities within each city. (c, d) Univariate relationships between phenological temperature sensitivity and city-level climate for the 10 most common species across cities. Points and vertical bars indicate species-specific estimates of phenological sensitivity to temperature *β*_T_ with 95% credible intervals. Solid lines show fitted relationships between phenological temperature sensitivity and individual climatic variables, estimated by repeatedly regressing posterior draws of the temperature coefficient on either city-level MAT or MAP. Shaded areas represent the 95% uncertainty bands across repeated fits.

For EOS, the average response across species was positive in 54 of 75 cities (Fig. 4b). Compared with SOS, EOS showed more widespread responses to urban warming, consistent with the overall species–city patterns shown in Fig. 2. Nevertheless, the direction of the EOS response was not uniform across all cities. In several hotter or drier cities, warmer climatic conditions were instead associated with earlier fall senescence. For example, when averaged across species within each city, a 1 ^◦^C increase in fine-scale temperature was associated with advanced EOS by 4.51 days in St. Augustine, FL, 3.40 days in Austin, TX, 2.97 days in Sacramento, CA, and 2.68 days in San Jose, CA. The mean intra-city among-species SD was 3.26 days ^◦^C^−1^, with an interquartile range of 2.24–3.96 days ^◦^C^−1^, indicating that species’ fall responses also differed substantially within cities, with stronger among-species variance than observed for SOS.

Among the 10 most common species, phenological temperature sensitivities showed clear relationships with city-level climate (Fig. 4c, d). For SOS, higher fine-scale temperatures tended to advance leaf-out for species planted in colder cities, but delay SOS for species growing in hot cities, with the crossover point occurring at around 10 ^◦^C of mean annual temperature (bio1; Fig. 4c). In contrast, higher intra-city temperatures tended to delay the EOS for species in wetter cities, but advance EOS in drier cities, with the crossover point occurring at around 1,000 mm of mean annual precipitation (bio12; Fig. 4d).

Multivariate models across all species showed similar climate-dependent patterns (Fig. 5). For SOS, temperature sensitivity was negatively associated with mean annual precipitation (mean = -0.10, 95% credible interval [−0.15, −0.05]) and positively associated with mean annual temperature (mean = 0.17, 95% credible interval [0.11, 0.23]), suggesting moisture and thermal constraints on spring phenology (Fig. 5a). For EOS, temperature sensitivity showed a strong positive relationship with mean annual precipitation (mean = 0.68, 95% credible interval [0.60, 0.75]), suggesting that warmer intra-city temperature delayed senescence in wetter cities but advanced senescence in drier cities across our study area (Fig. 5b).

**Figure 5:**
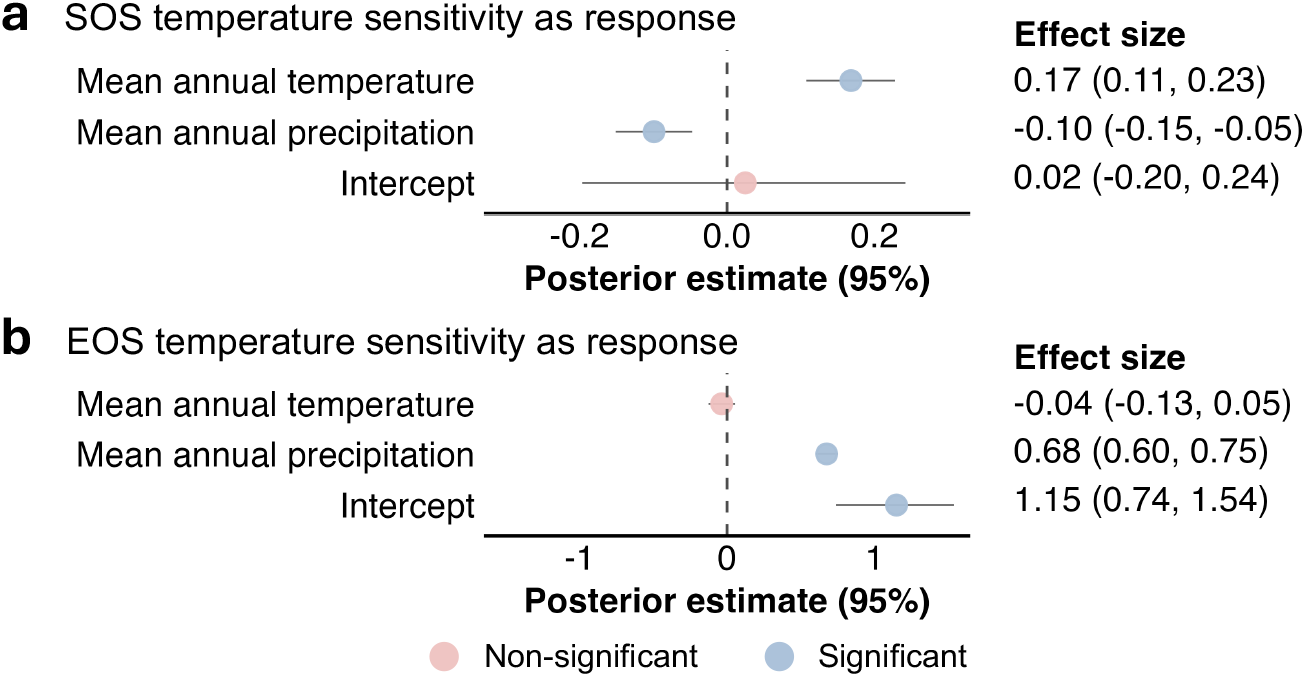
Relationships between phenological temperature sensitivity and city-level climate. Fixed-effect estimates from phylogenetic generalized linear mixed models with SOS temperature sensitivity (a) and EOS temperature sensitivity (b) as response variables. Points indicate posterior mean estimates, and horizontal lines show 95% credible intervals.

## 4 Discussion

In this study, we integrated municipal street-tree inventories, PlanetScope-derived phenology, and 1-km near-surface urban air-temperature data to examine individual-tree phenology across 76 cities in the contiguous United States. Our results support three main conclusions. First, phenological variation within cities was often comparable to variation among cities for the same species, indicating that urban tree phenology cannot be fully represented by city-level averages. Second, phenological timing and growing season length were widely associated with fine-scale intra-city temperature variation, although the direction and magnitude of these associations differed among species, cities, and phenological phases. Third, phenological sensitivity to temperature varied systematically with regional climatic context: warmer local urban environments were generally associated with earlier spring onset and delayed fall senescence in cooler or wetter cities, whereas these associations weakened or reversed in hotter or drier cities. Together, these findings show that urban tree phenology reflects interactions among local thermal heterogeneity, species identity, and regional climate, with implications for urban forest monitoring, species selection, and climate adaptation.

### 4.1 Fine-scale urban temperature and intra-city variation in phenology

Our results show that urban phenology cannot be treated as a uniform city-level average (Questions 1 and 2). For many species, individual-tree phenological variation within cities was comparable to, or larger than, variation among cities (Fig. 1 and Fig. 2). This finding is consistent with previous city-specific case studies in the U.S., showing that temperature regulates urban phenology through local micrometeorological conditions, despite differences in data sources (Crawford et al., 2024; Katz et al., 2019; Parece and Campbell, 2018; Wu et al., 2025; Zipper et al., 2016). Our study advances this literature by revealing the broader generality of phenological heterogeneity and fine-scale temperature sensitivity across species and cities: these patterns are evident across a broad set of urban tree inventory species, and remain apparent across cities spanning diverse climatic regions (Fig. 2 and Fig. 3). The individual-level framework captures species-specific and within-city spatial details that are often unresolved by coarser-resolution sensors (Badeck et al., 2004; Zeng et al., 2020; Zhao et al., 2022).

These results also highlight the importance of fine-scale urban thermal heterogeneity. Previous studies have often examined urban phenology through urban–rural contrasts, macroclimatic gradients, or proxies for urbanization such as impervious surface fraction, land surface temperature, or population density (Dallimer et al., 2016; Jia et al., 2021; Li et al., 2017; Meng et al., 2020; Wang et al., 2025, 2019). By contrast, our analysis links individual-tree phenology to near-surface air temperature within cities, allowing us to evaluate phenological sensitivity to temperature across many species–city combinations. Because air temperature is a direct environmental cue for plant phenology, this approach provides a closer representation of the thermal conditions relevant to trees than many indirect urbanization metrics (Chakraborty et al., 2022; Du et al., 2021).

The observed intra-city heterogeneity has practical implications for urban forest management. In many species–city combinations, earlier SOS and later EOS under warmer local conditions were accompanied by longer GSL (Fig. 2b, d), indicating that fine-scale urban thermal variation can alter not only the timing of individual phenological events but also the duration of seasonal canopy activity. By altering the timing and magnitude of canopy-mediated functions, particularly seasonal carbon uptake and release, variation in individual-tree phenology may scale up to neighborhood-level differences in ecosystem services and disservices, including cooling benefits and pollen exposure (Crawford et al., 2024; Katz et al., 2019; Song et al., 2025). Management strategies based only on citywide averages may overlook these fine-scale differences. Incorporating high-resolution monitoring of both urban thermal environments and tree phenology could improve urban forest planning under continued warming (Hall et al., 2016).

### 4.2 Regional climate modulates phenological sensitivity to urban temperature

Phenological sensitivity to temperature varied not only among species and cities, but also across regional climatic gradients. Spring phenology showed weaker and more variable associations with local temperature than fall phenology, whereas fall senescence was more consistently delayed in warmer local urban environments (Fig. 2, Fig. 4a, b). This difference between SOS and EOS is consistent with broader phenological literature showing that spring and fall phenology are regulated by different combinations of environmental cues and physiological constraints. Spring leaf-out depends on interactions among winter chilling, spring forcing, photoperiod, and species-specific requirements (Ettinger et al., 2020; Fu et al., 2015; Hänninen et al., 2019; Li et al., 2022; Meng et al., 2020), whereas fall senescence can be strongly influenced by late-season temperature, water availability, carbon balance, and stress events (Bigler and Vitasse, 2021; Chaves, 1991; Fu et al., 2018; Luo et al., 2024).

The climate-dependent patterns observed here suggest that local warming within cities does not have uniform phenological impacts. In cooler cities, warmer local urban environments were generally associated with earlier spring onset. In hotter cities, however, this association weakened or became positive, indicating delayed SOS in warmer local environments (Fig. 2a, Fig. 4a, c). This pattern is consistent with the possibility that additional warming in already warm environments may reduce chilling accumulation or push trees closer to thermal stress thresholds, thereby limiting or reversing spring advancement (Liang et al., 2024; Yin et al., 2024). However, because we did not directly quantify chilling fulfillment, forcing accumulation, or tree physiological status, this mechanism needs a direct test in future studies.

For fall phenology, warmer local urban environments were generally associated with delayed senescence in wetter cities but earlier senescence in drier cities (Fig. 2b, Fig. 4b, d). This pattern is consistent with moisture limitation modifying the effect of local warming on fall phenology. In wetter environments, warming may extend photosynthetic activity and delay senescence when water limitation is weak. In drier environments, however, warmer local conditions may increase evaporative demand, intensify drought stress, or coincide with more impervious and sparsely vegetated microsites, leading to earlier senescence (Franceschi et al., 2023; Liang et al., 2024; Stratópoulos et al., 2019; Yin et al., 2024). Because soil moisture, vapor pressure deficit, and irrigation were not directly measured, these explanations should be viewed as plausible mechanisms for future testing.

These findings complement previous studies showing that urban phenological responses depend on regional climate (Li et al., 2019; Meng et al., 2020; Yang et al., 2023). However, our results extend this perspective from urban–rural or city-level contrasts to individual trees within cities. The same degree of local urban warming may be associated with different phenological outcomes depending on whether a city is cool or warm, wet or dry. As a result, models that assume uniform phenological responses to urban warming may misestimate growing-season length, canopy function, and species suitability in urban forests (Fu et al., 2023; Li et al., 2019).

### 4.3 Species-specific phenological sensitivity and its potential drivers

Species identity was an important axis of phenological heterogeneity for urban trees (Figs. 2 and 4a,b). Even within the same city, co-occurring species often differed in both phenological timing and phenological sensitivity to temperature. Similarly, the same species sometimes showed different temperature sensitivities across cities. These patterns indicate that urban phenology reflects not only local environmental conditions, but also the taxonomic composition of planted urban forests. Species-specific responses are expected because tree species differ in phenological cues, chilling and forcing requirements, drought tolerance, hydraulic strategies, photoperiod sensitivity, and stress tolerance (Cole and Sheldon, 2017; Wu et al., 2025). In urban settings, these intrinsic differences may be amplified by planting practices. Many urban tree species are planted outside their native climatic ranges or in environments that differ from those experienced by source populations (Chamberlain and Wolkovich, 2023; Ghelardini et al., 2006; Jensen et al., 2022; Linkosalo and Lechowicz, 2006). Provenance and nursery origin may therefore influence phenological responses, because phenology is often heritable and locally adapted (Chamberlain and Wolkovich, 2023; Iler et al., 2021; Wu et al., 2025). For example, southern-origin populations often show delayed budburst in cooler or higher-latitude urban environments because of differences in photoperiod sensitivity, chilling requirements, and temperature thresholds (Ghelardini et al., 2006; Jensen et al., 2022; Linkosalo and Lechowicz, 2006), while northern provenances may require fewer growing degree days and thus respond more readily to urban warming (Chamberlain and Wolkovich, 2023). Future work incorporating detailed climatic legacy, planting history, and physiological processes will be needed to disentangle these effects.

These species-level differences have important implications for urban forest planning. As cities warm, species that maintain appropriate phenological timing under local thermal and moisture conditions may be better able to sustain canopy function, cooling, carbon uptake, and biotic interactions (Dallimer et al., 2016; Ettinger et al., 2022; Harrison and Winfree, 2015). Conversely, species showing delayed spring onset or earlier fall senescence in warmer or drier local environments may be more vulnerable to functional stress, especially if these phenological shifts reflect unmet chilling requirements, heat stress, or water limitation in urban environments (Du et al., 2022; Zani et al., 2020). Phenological sensitivity to temperature may therefore provide an early indicator of species performance under urban climate stress, complementing occurrence-based or demographic assessments of urban tree suitability (Burley et al., 2019; Esperon-Rodriguez et al., 2022; Wu et al., 2025; Zhu et al., 2025).

### 4.4 Limitations and future directions

Several limitations should be considered when interpreting our results. First, although PlanetScope provides frequent observations at relatively high spatial resolution, satellite-derived phenology for individual urban trees remains uncertain. Extracted EVI time series may include signals from neighboring crowns, understory vegetation, turfgrass, impervious surfaces, or other nearby land-cover features. This issue is likely more pronounced for small trees, such as those with canopy diameters smaller than 3 m, whose crowns may not be fully resolved at PlanetScope resolution (Song et al., 2025; Wu et al., 2021). Our validation against NPN observations, together with previous evaluation analyses, showed that PlanetScope-derived phenology captured meaningful seasonal signals, but agreement differed between SOS and EOS (Song et al., 2025; Zhao et al., 2022). Future work using standardized field observations, drone imagery, or high-resolution aerial data could better quantify uncertainty in individual-tree phenology retrieval.

Second, municipal tree inventories differ in timing, completeness, taxonomic resolution, and location accuracy (McCoy et al., 2022). Some inventories may predate the PlanetScope observation period, and tree removals, new plantings, or changes in health condition may not be fully captured. Species misidentification or coordinate error could also contribute to apparent intra-city phenological variation. In addition, our inferences apply primarily to inventoried deciduous street trees, not all urban trees. Park trees, private-yard trees, and remnant urban forest patches may show different phenological patterns because they differ in canopy structure, soil conditions, management, and disturbance history.

Third, although U-HAT provides high-resolution near-surface air temperature across cities, its 1-km resolution does not capture all tree-scale thermal variation. Street trees can experience substantial microclimatic differences at meter-to tens-of-meters scales due to shading, building geometry, pavement, traffic, irrigation, and local canopy cover. Therefore, our analysis characterizes fine-scale intra-urban temperature variation at the neighborhood scale rather than the full microclimatic environment experienced by each tree. Integrating sub-kilometer air-temperature observations, mobile sensing, or sensor networks would improve the ability to estimate tree-scale thermal exposure.

Fourth, our seasonal temperature metrics summarize broad preseason conditions and may not fully capture the phenological cues that regulate SOS and EOS. Spring phenology is shaped by the interaction between winter chilling and spring forcing, while fall phenology may respond to late-season heat, cold spells, drought stress, and short-term weather variability (Chuine et al., 2010; Fu et al., 2015; Wang et al., 2020). Future analyses using chilling accumulation, growing degree days, vapor pressure deficit, soil moisture, and event-based temperature metrics could improve mechanistic attribution.

Finally, other urban environmental factors may contribute to intra-city phenological heterogeneity. Artificial light at night, in particular, can delay fall senescence and modify phenological sensitivity to temperature and precipitation (Chen et al., 2026; Friulla and Varone, 2025; Meng et al., 2022; Wang et al., 2025). Land-use composition, irrigation, soil compaction, pollution, and management intensity may also interact with temperature to influence phenological timing. Integrating these variables with individual-tree phenology and urban climate data would provide a more complete understanding of the environmental drivers of urban phenological variation.

### 4.5 Conclusion

In this study, we showed widespread intra-city phenological variation and fine-scale temperature-phenology associations across 76 cities in the contiguous United States. Furthermore, we found the magnitude and direction of these associations varied substantially among species, and were modulated by regional climatic conditions. Recognizing that urban tree phenology emerges from interactions among local thermal heterogeneity, species identity, and regional climate can support both fine-scale monitoring of phenological responses and more informed species selection under continued urban warming. Our findings also suggest the need to move beyond single-driver explanations and more fully investigate the complex, context-dependent processes that shape urban tree phenology across spatial scales.

## Acknowledgments

We thank Dr. Karen Seto and members of the Zhu Lab at the University of Michigan for providing helpful advice at earlier stages of this work. Jiali Zhu was supported by the Institute for Global Change Biology (IGCB) Graduate Fellowship at the University of Michigan. Kai Zhu and Jiali Zhu were supported by the National Science Foundation [grant numbers 2306198 (CAREER)]

## Competing interests

There are no competing interests to declare.

## Notes

### Competing Interest Statement

The authors have declared no competing interest.

